# Cytoplasmic calcium influx mediated by Lr14a regulates stomatal immunity against leaf rust in wheat

**DOI:** 10.1101/2024.12.10.627212

**Authors:** Lili Yue, Limin Wang, Benjamin Neuhäuser, Songyuan Zhang, Gerhard Herren, Matthias Heuberger, Esther Jung, Uwe Ludewig, Cyril Zipfel, Beat Keller

**Author notes:** Corresponding authors Correspondence and requests for materials should be addressed to L.Y., C.Z. and B.K.

## Abstract

The race-specific resistance gene *Lr14a* in wheat confers a unique type of heterogenous resistance reaction. It encodes an ankyrin-repeat transmembrane domain protein that confers immunity against the fungal pathogen *Puccinia triticina*. Here, we show that Lr14a functions as a calcium-permeable channel, mediating cytoplasmic Ca²⁺ influx that is crucial for leaf rust resistance in wheat. Infection with avirulent isolates induced *Lr14a* expression predominantly in mesophyll cells while triggering cell death in guard cells in wheat. This study revealed a mechanism by which the product of an *R* gene regulates stomatal immunity non-cell autonomously through the mediation of calcium signaling.

---

Leaf rust, caused by the fungal pathogen *Puccinia triticina*, poses a serious threat to wheat production worldwide^1^. Unlike other fungal pathogens, *P. triticina* specifically recognizes stomata as entry points for appressorium formation and subsequent leaf invasion^2^. Studies have shown that stomatal closure and guard cell death are critical steps in wheat resistance (*R)* gene-mediated, race-specific resistance to fungal pathogens^3–6^. Despite these findings, the mechanisms by which *R* genes regulate stomatal immunity have remained largely unexplored.

In plant-pathogen interactions, *R* genes often encode immune receptors, such as nucleotide-binding domain, leucine-rich repeat (NLRs) proteins, helper NLRs, receptor-like proteins (RLPs), and receptor-like kinase (RLKs), which detect pathogen effectors and frequently trigger hypersensitive response (HR)-associated host cell death, restricting pathogen growth^7^. Recent research has shown that *Arabidopsis* NLRs, helper NLRs, and mixed-lineage kinase domain-like proteins (MLKLs) can function as Ca²⁺ channels or mediate Ca²⁺ influx to trigger host cell death^8–10^. Similarly, in wheat, the NLR encoded by the stem rust resistance gene *Sr35* forms a complex with the AvrSr35 effector, resulting in a resistosome that functions as a Ca²⁺-permeable non-selective cation channel, inducing host cell death^11^.

*Lr14a* confers a unique, mesothetic (heterogenous) resistance phenotype with different infection types on the same leaf^12^. *Lr14a* alone is no longer an effective resistance gene in wheat production as virulent races are widely present^12^. Interestingly, *Lr14a* is one of three major quantitative trait loci which together confer durable, broad-spectrum resistance in the wheat cultivar Forno^13,14^: in combination with the ABCG transporter encoded by *Lr34*^15^ and the *Lr75* gene^16^, *Lr14a* confers broad-spectrum and durable resistance, making it a highly valuable gene in gene combinations.

*Lr14a* encodes a protein containing an ankyrin-repeat transmembrane (ANK-TM) domain with sequence similarities to two known calcium channels: *Arabidopsis* ACD6 and human TRPA1^17,18^. Our previous research demonstrated that *Lr14a* expression is low or undetectable in uninfected plants but significantly upregulated following infection with the avirulent *P. triticina* isolate 96209, peaking at four days post-infection (dpi). Transient expression of *Lr14a* in *Nicotiana benthamiana* induced a water-soaking-like phenotype, which was inhibited by the calcium channel blocker lanthanum (III) chloride (LaCl₃)^13^. These characteristics imply that Lr14a has a unique function in plant immunity. Further characterization of Lr14a has the potential to reveal fundamentally novel molecular mechanisms on how plants achieve race-specific immunity.

In this study, we characterized Ca^2+^ conductivity of Lr14a using both heterologous systems and wheat. The Lr14a-mediated Ca^2+^ influx is essential for leaf rust resistance. Lr14a specifically induced guard cell death in wheat and *N. benthamiana*. However, when *Lr14a* was expressed in wheat protoplasts or epidermal cells, no cell death was observed, implying that cell death in guard cells is non-cell autonomous.

## Results

### Lr14a is a calcium-permeable channel

Lr14a shares structural similarity with the transient receptor potential cation channel TRPA1, and the Lr14a-induced water-soaking-like phenotype can be abolished by the Ca²⁺ influx channel blocker LaCl_3_ ^13,18^. These observations prompted us to investigate whether Lr14a functions as a Ca²⁺-permeable channel. To test this, we first measured Ca²⁺ influx in transgenic *N. benthamiana* carrying the Ca²⁺ reporter GCaMP3^19^ after transient expression of *Lr14a* and its two loss-of-function mutants, *L362F* and *G768S^13^.* A β-estradiol-inducible promoter was used to induce gene expression. The protein accumulation levels were comparable after β-estradiol treatment (Extended Data Fig. 1a). We observed sustained cytoplasmic Ca^2+^ influx in Lr14a-expressing leaves, whereas the two loss-of-function mutations L362F and G768S significantly reduced Lr14a-mediated cytoplasmic Ca^2+^ influx (Fig. 1a) and did not show the water-soaking-like phenotype (Extended Data Fig. 1d). Furthermore, the general Ca^2+^ influx channel blockers LaCl₃ and GdCl₃ abolished both the Lr14a-mediated water-soaking-like phenotype and cytoplasmic Ca^2+^ influx (Extended Data Fig. 1e, f and Fig. 1b, c), suggesting that Lr14a facilitates Ca²⁺ transport in *N. benthamiana*, and amino acids L362 and G768 were essential for Ca²⁺ transport activity.

**Figure. 1.**
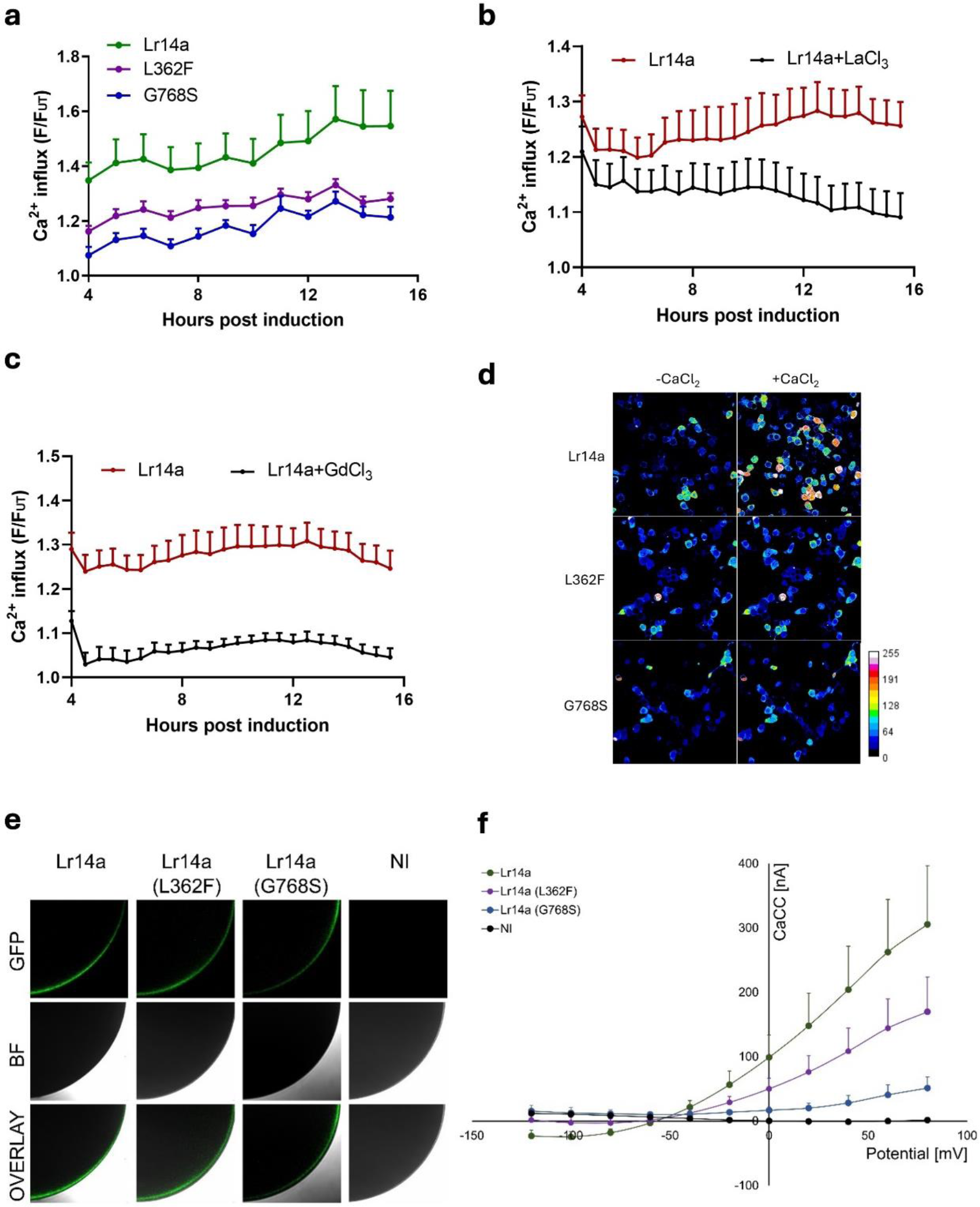
Lr14 forms an ion channel permeable to Ca^2+^. **a-c**, Time-course measurements of Lr14a-induced Ca^2+^ influx in transgenic *N. benthamiana* leaves expressing GCaMP3. Protein expression was induced by 100 μM β-estradiol. Two loss-of-function mutants, L362F and G768S, show reductions in Ca^2+^ influx (**a**). Lr14a-induced Ca^2+^ influx was abolished by the application of LaCl₃ (**b**) and GdCl₃ (**c**). Data were collected from 15-18 individual leaves from three independent experiments, shown as mean ± SE. Average Ca^2+^ influx was calculated in Extended Data Fig. 1g-i. **d**, Increases in [Ca^2+^]_cyt_ in response to 5mM Ca^2+^ in HEK293 cells expressing flag-tagged Lr14a, or its mutants L362F and G768S. Changes in [Ca²⁺]_cyt_ were visualized using GCaMP6m and scaled with a pseudo-color bar. Protein accumulation was determined by western blot shown in Extended Data Fig. 2b). **e**, Localization of Lr14a-GFP fusion protein variants in oocytes. Lr14a–GFP, L362F-GFP, G768S-GFP fusion protein localization in the plasma membrane. Control shows non-injected oocytes (NI). Dark images show GFP fluorescence in the membrane, bright image shows the bright-field image of the oocyte and the lowest image shows the overlay of both. One representative picture of n=20 is shown. **f**, Ca^2+^ induced chloride currents (CaCC) in oocytes expressing Lr14a and its variants. Current / Voltage plot showing currents induced by 10 mM CaCl_2_ in dependence of the membrane potential. Lr14a is shown as closed green circles. L362F and G768S are shown as closed purple and blue circles, respectively. Non-injected controls are shown as black circles. All currents were recorded at pH 5.5 from the same batch of oocytes. Data is given as means ± SE (n=4).

To determine if Lr14a directly mediates Ca²⁺ influx, we transiently expressed Lr14a with calcium indicator protein GCaMP6m in human embryonic kidney 293 (HEK293) cells following subsequent extracellular Ca²⁺ treatment. The protein accumulation levels were comparable (Extended Data Fig. 2b). The increase of extracellular Ca²⁺ concentration from 0.1 mM to 5 mM induced a sustained [Ca^2+^]_cyt_ increase in Lr14a-expressing cells. The loss-of-function mutations L362F and G768S significantly reduced Lr14a-mediated Ca²⁺ influx (Fig. 1d and Extended Data Fig. 2a), which is consistent with the observation in *N. benthamiana* (Fig. 1a).

Finally, an electrophysiological experiment was carried out to determine Ca²⁺ channel activity of Lr14a. We measured Ca²⁺-induced chloride currents (CaCC) in *Xenopus* oocytes, as direct observation of calcium influx is hindered by the presence of endogenous calcium-activated chloride channels. Lr14a localized to the plasma membrane and induced CaCC in oocytes, while L362F and G768S exhibited reduced CaCC compared to wild type Lr14a (Fig. 1e, f).

We then asked whether Lr14a induces a calcium signal in wheat. In our previous work^13^, *Lr14a* expression was very low or undetectable in uninfected plants but increased after infection with the avirulent *P. triticina* isolate 96209, reaching peak expression at 4 days post-infection (dpi), which is not observed after infection with a virulent isolate 95037. Thus, we measured Ca²⁺ flux in the Arina*LrFor* (*Lr14a* near-isogenic line [NIL]) and two Arina*LrFor*-derived EMS loss-of-function susceptible mutants at 4 dpi with avirulent isolate 96209 using the Non-invasive Micro-test Technology (NMT). After infection, the wild-type Arina*LrFor* exhibited significantly higher sustained Ca²⁺ influx compared to the two loss-of-function susceptible mutants L362F and W680* (early truncation mutation) (Fig. 2a-c). We also analyzed Ca²⁺ flux before and after infection. The avirulent isolate 96209-infected line exhibited higher Ca²⁺ influx compared to the mock treatment (Fig. 2d, e), while the virulent isolate 95037-infected line showed no significant difference compared with the mock (Extended Data Fig. 3a, b). These results suggest that avirulent isolate-induced *Lr14a* expression triggers Ca²⁺ influx in wheat. Collectively, these results demonstrate that Lr14a forms a Ca²⁺-permeable channel mediating cytoplasmic Ca^2+^ influx, which is essential for Lr14a-based leaf rust resistance.

**Figure. 2.**
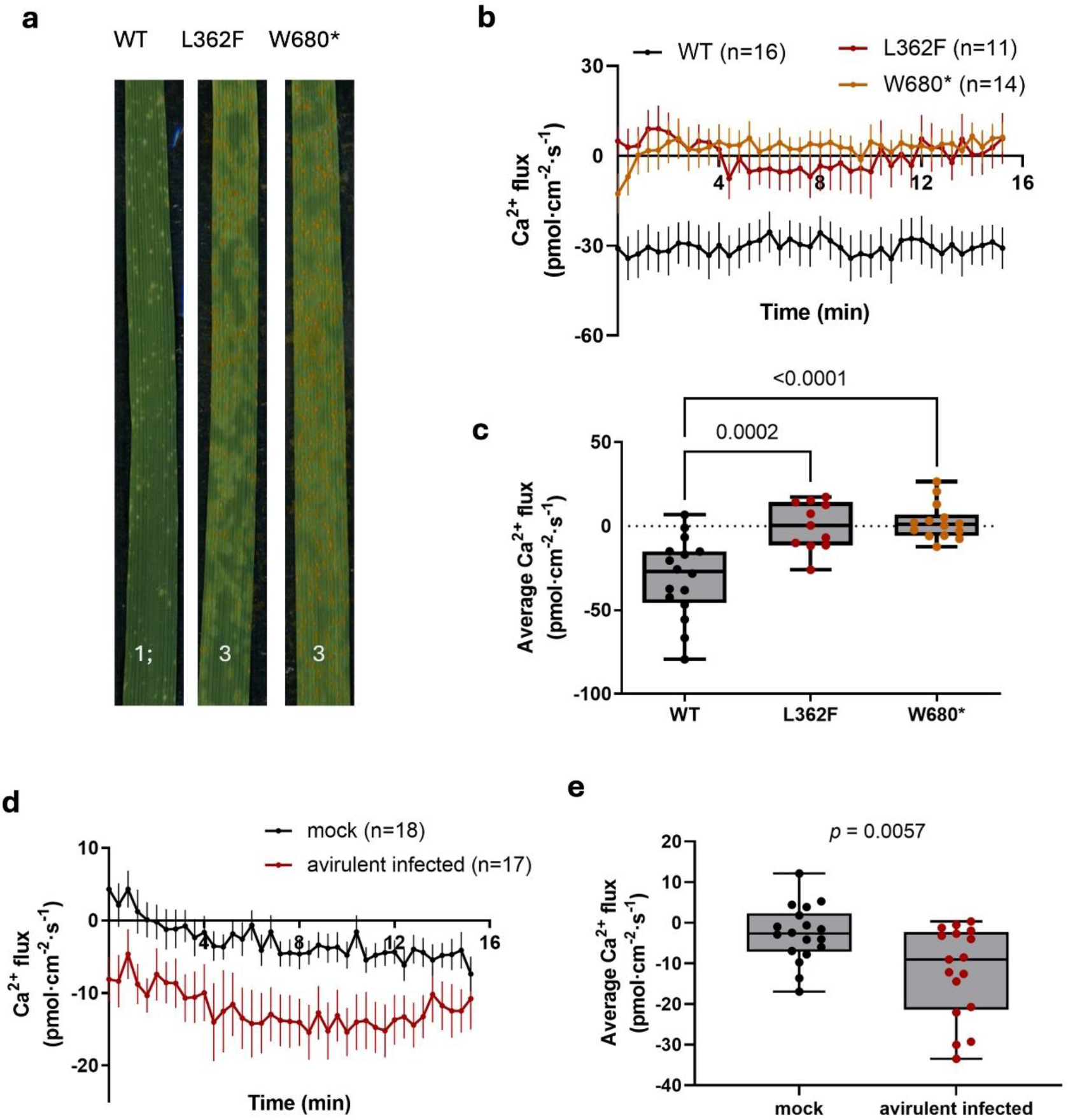
Lr14a mediates Ca^2+^ influx in wheat. **a**, Seedling infection of Arina*LrFor* (WT) and two Arina*LrFor* -derived loss-of-function mutants with avirulent *P. triticina* isolate 96209. 1 = small uredia with necrosis, 3 = medium sized uredia with or without chlorosis,; = hypersensitive flecks. **b, d**, Time-course recordings of Ca^2+^ flux in mesophyll cells of wheat using NMT. Negative values indicate Ca^2+^ influx, and positive value indicate Ca^2+^ efflux. **b**, Ca^2+^ flux in Arina*LrFor* (WT) and two loss-of-function mutants. **c** Quantitative analysis of average Ca^2+^ fluxes from (**b**). **d**, Ca^2+^ flux in Arina*LrFor* (WT) infected with avirulent isolate 96209 and mock. **e**, Quantitative analysis of average Ca^2+^ fluxes from (**d**). All the data are presented as mean ± SE. Statistical significance was determined using an unpaired, two-tailed *t*-test.

### Lr14a induces guard cell death in *Nicotiana benthamiana*

Our previous study indicated that Lr14a induces a water-soaking-like phenotype in *N. benthamiana*. It has been described in bacterial pathosystems that the water-soaking phenotype is associated with pathogen effectors, since high humidity in the apoplast facilitates pathogen multiplication^20,21^. Therefore, we hypothesized that the observed Lr14a-triggered water-soaking-like phenotype might not be classical water-soaking but rather cell death, since *R* genes trigger host cell death to inhibit pathogen growth.

To test this, we performed a trypan blue-based staining for dead cells with *N. benthamiana* leaves expressing HA-tagged Lr14a and two loss-of-function mutants, L362F and W680*. The ADR1 was used as positive control. Trypan blue staining revealed that leaves expressing Lr14a underwent cell death in *N. benthamiana*, whereas the L362F and W680* mutants showed no cell death-inducing activity (Extended Data Fig. 4a). However, Lr14a-induced cell death was significantly weaker than that caused by ADR1. Additionally, the cell death was only visible on abaxial leaf surface, even 10 days post-infiltration (Extended Data Fig. 4a, b), leading us to hypothesize that cell death may be restricted to the abaxial epidermis.

To test this hypothesis, we peeled the abaxial epidermis of *N. benthamiana* leaves three days after infiltration with Lr14a or its mutants, L362F and W680*. We then treated the epidermis with fluorescein diacetate (FDA), which specifically stains living cells. We observed that guard cells in leaves expressing Lr14a underwent cell death, as indicated by the absence of an FDA signal (Extended Data Fig. 4c). In contrast, guard cells expressing the L362F and W680* mutants remained viable (Extended Data Fig. 4c). FDA-stained epidermal cells were not observed in any of the datasets, likely because guard cells are more metabolically active and exhibit higher esterase activity than other epidermal cells^22^. A five-minute incubation with FDA appears sufficient for staining guard cells but may be inadequate for other epidermal cell types. We then isolated protoplasts of *N. benthamiana* expressing Lr14a and its mutants and assessed their viability. All leaves contained healthy protoplasts, regardless of whether they expressed Lr14a, L362F, or W680* (Extended Data Fig. 5a).

We then performed FDA and propidium iodide (PI, stains dead cells) staining on *N. benthamiana* intact leaves. PI primarily stained guard cells of the leaves expressing Lr14a, while no PI signal was detected in those expressing L362F and W680* (Extended Data Fig. 5b). In contrast, FDA staining revealed that both guard cells and surrounding epidermal cells were stained in leaves expressing L362F or W680*, whereas only the epidermal cells were stained in Lr14a-expressing leaves, with guard cells remaining unstained (Extended Data Fig. 5c). These findings suggest that Lr14a specifically induces guard cell death in *N. benthamiana*, potentially explaining the Lr14a-triggered water-soaking-like phenotype as a consequence of partial cell death in the guard cells of the abaxial epidermis.

### Lr14a induces both stomatal closure and guard cell death in wheat

Based on our observations in *N. benthamiana* and considering the important role of stomata in *P. triticina* penetration, we investigated Lr14-mediated stomatal behavior in Arina*LrFor* near-isogenic line (NIL) expressing Lr14a. Stomatal apertures were measured at 4 dpi, coinciding with the peak expression of *Lr14a*, with avirulent isolate 96209 and virulent isolates 95037. The abaxial epidermis of infected wheat leaves were peeled, and the stomatal apertures were quantified microscopically. The results showed that the avirulent isolate 96209 induced stomatal closure in wheat, which was significantly stronger than the induction by the virulent isolate 95037 (Fig. 3a).

**Figure. 3.**
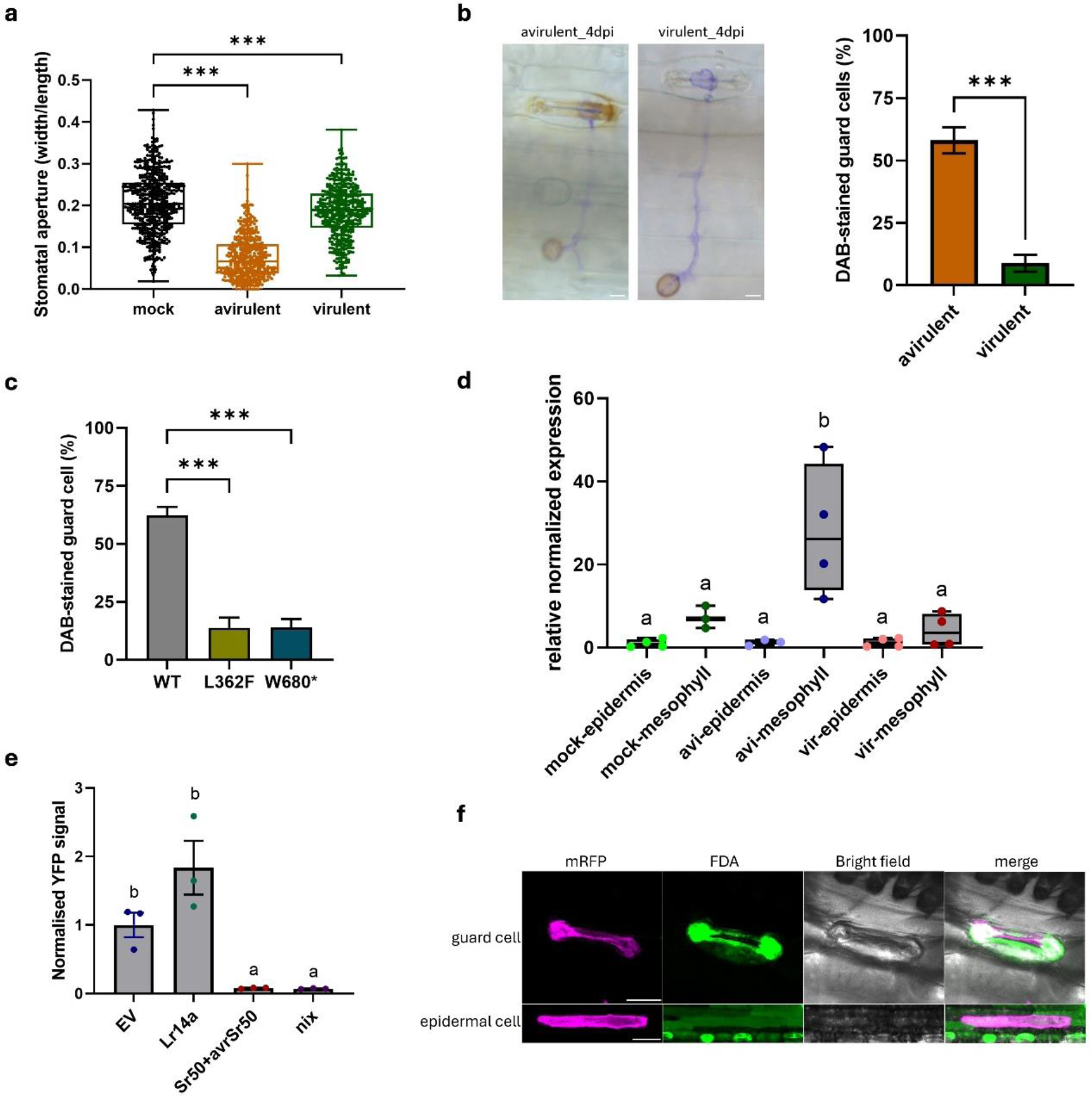
Lr14a triggers guard cell death in wheat. **a**, Stomatal aperture measurement of Arina*LrFor* infected with avirulent isolate 96209 and virulent isolate 95037 at 4dpi. Data represents more than 450 stomata from three independent experiments. **b**, DAB-stained guard cells in Arina*LrFor* challenged with avirulent isolate 96209 and virulent isolate 95037 at 4dpi revealed by DAB staining. Left: representative picture of appressorium-formed stomata after DAB staining. Scale bar = 30 μm. Right: quantitative analysis of DAB-stained guard cells among appressorium-formed stomata. n > 112 guard cells from two independent experiments. **c**, Comparison of DAB-stained guard cell in wild-type (WT) Arina*LrFor* and loss-of-function mutants (L362F and W680* mutants) after infection with avirulent isolate 96209 at 4 dpi. The percentage of dead guard cells, as indicated by DAB staining, was calculated by considering only stomata with a visible appressorium. n > 83 guard cells from two independent experiments. **d**, *Lr14a* expression in epidermis and mesophyll cell after infection with avirulent isolate 96209 (avi) and virulent isolate 95037 (vir) at 4dpi. n = 4, with different letters indicating statistical significance (*P* < 0.05). **e**, Protoplasts of the wheat cultivar Bobwhite S26 were co-transformed with YFP and either empty vector (EV), Lr14a driven by a ubiquitin promoter, or Sr50/avrSr50 (positive control). YFP signal from EV-transformed protoplasts was normalized to 1. Non-transformed protoplasts (nix) served as background control. n = 3, with different letters indicating statistical significance (*P* < 0.05). **f**, Epidermal cells of wheat cultivar Bobwhite S26 co-expressing *Lr14a* and *mRFP.* Particle bombardment was performed for transient expression. FDA staining was applied to assess cell viability. Scale bar 25 μm (upper) and 100 μm (lower). \*\*\**P* < 0.0001 (ordinary one-way ANOVA with multiple comparisons; unpaired *t*-test with two-tailed for two-group comparisons).

3,3’-diaminobenzidine (DAB) staining was performed to assess the hypersensitive response (HR) of Arina*LrFor* after infection with avirulent and virulent isolates at 4 dpi. We quantified the percentage of DAB-stained guard cells specifically within stomata where an appressorium had formed. In leaves infected with the avirulent isolate 96209, 58 % of the guard cells were stained by DAB, which was significantly higher compared to 9 % with the virulent isolate 95037 (Fig. 3b). Additionally, two Arina*LrFor*-derived loss-of-function mutants, L362F and W680*, exhibited reduced DAB-stained guard cells compared to wild-type Arina*LrFor* after infection with the avirulent isolate 96209 at 4 dpi (Fig. 3c). These results suggested that impairing the stomata function by inducing stomatal closure and guard cell death is a key step of Lr14a-mediated resistance.

### The expression of *Lr14a* is exclusively induced in mesophyll cells but triggers cell death in guard cells

Given that *Lr14a* expression in both *N. benthamiana* and wheat specifically induced guard cell death, we aimed to determine whether this phenotype is associated with a guard cell-specific expression of the *Lr14a* gene. To investigate this, we analyzed *Lr14a* expression in both epidermal and mesophyll cells of Arina*LrFor* using qRT-PCR after infection with avirulent and virulent *P. triticina* isolates at 4 dpi. The qPCR results showed that the avirulent isolate 96209 predominantly induced *Lr14a* expression in mesophyll cells, but not in epidermal cells. The *Lr14a* expression was low in both avirulent isolate 96209 and virulent isolate 95037 infected epidermis. Infection with the virulent isolate 95037 did not induce *Lr14a* expression in either mesophyll or epidermal cells. (Fig. 4d). The gene *TaALMT12*, which is specifically expressed in guard cells^23^, was used as control (Extended Data Fig. 6).

To further investigate whether *Lr14a* expression in guard cells has a direct association with the observed cell death, we performed single-cell transformation of wheat leaf epidermal cells in ‘Bobwhite S26’ using particle bombardment. *mRFP* was used as a reporter gene co-expressed with the target genes to identify transformed cells. FDA was used as an indicator of live cells. We observed that cells expressing the reporter *mRFP* were also stained with FDA, indicating that no cell death occurred in either guard cells or other epidermal cells after *Lr14a* expression, suggesting that the Lr14a-triggered guard cell death does not depend on an epidermal expression of *Lr14a* (Fig. 3f).

The inducible expression profile of *Lr14a*, its structural characteristics, including predicted transmembrane helices, and its ability to induce cell death, are reminiscent of TALE-activated executor resistance genes^24^. To determine whether *Lr14a* expression in mesophyll cells is sufficient to trigger cell death as an executor *R* gene, we conducted a protoplast cell death assay in the wheat cultivar ‘Bobwhite S26’. Yellow fluorescent protein (YFP) was used as a reporter gene co-expressed with the target genes to indicate cell viability. In protoplasts co-transfected with the positive control *Sr50* and *AvrSr50*, a strong reduction of YFP signal was observed, indicating cell death was triggered. In contrast, no reduction in YFP signal was detected in protoplasts transfected with *Lr14a*, with a comparable signal to protoplasts transformed with the empty vector (EV). This suggests that *Lr14a* expression did not trigger cell death in protoplasts (Fig. 3e).

These results indicated that guard cell death is not a consequence of specific *Lr14a* expression in guard cells. Instead, the exclusive induction of Lr14a in mesophyll cells may indirectly regulate guard cell activity via cell-to-cell communication.

## Discussion

Here, we found in four different experimental approaches and systems that Lr14a is a Ca²⁺ permeable channel. Expression of *Lr14a* in both *N. benthamiana* and wheat specifically triggered guard cell death. Notably, this effect did not rely on guard cell-specific expression. Instead, Lr14a expression was activated specifically in mesophyll cells in response to infection by an avirulent isolate. This activation appears to induce guard cell death non-cell autonomously.

*Lr14a* plays a critical role in leaf rust resistance. *Lr14a*, in combination with *Lr75* and *Lr34* (encoding an ABCG transporter) confers durable resistance against leaf rust in the winter wheat variety Forno^13,14,16,25^. Thus, despite the abundant occurrence of leaf rust isolates virulent on *Lr14a*, *Lr14a* contributes to durable adult plant resistance in the field. It is interesting to note that calcium-permeable cation channels encoded by NLRs when combined with the ABCG transporter Lr34 also result in durable, broad-spectrum resistance: Lr34 combined with the NLR Lr13^26^ confers durable leaf rust resistance^27^, whereas Lr34/Yr18 combined with Yr27 (an allelic variant of Lr13) confers durable stripe rust resistance^28^. Further biochemical studies are needed to understand the molecular control of this additive, non-race specific, durable resistance involving calcium ion and other substrate fluxes at the plasma membrane. It is tempting to speculate that durable resistance results from a combination of Lr34 ABC transport activity with a calcium-permeable cation channel.

In this study, we provide new insights into plant immunity by demonstrating that the race-specific *R* gene *Lr14a* encodes a calcium transporter that specifically triggers guard cell death, potentially preventing pathogen penetration. Although recent studies have shown that NLRs and helper NLRs can act as Ca²⁺ channels to initiate host cell death^8–10^, none of the previously identified *R* genes with calcium channel activity has been specifically associated with stomatal immunity in wheat. Interestingly, Lr14a triggers only partial guard cell death at a rate of 58% (Fig. 3b), which is consistent with the heterogenous leaf rust infection types observed in Lr14a containing plants^12^. We hypothesize that *P. triticina* can successfully penetrate through stomata at the early stage, releasing effectors into the mesophyll cells that activate Lr14a expression and initiate a resistance signaling pathway. This signaling subsequently induces guard cell death, disrupting later infections. The selective induction of guard cell death likely reflects the specialized recognition mechanisms between the stomata-targeting leaf rust pathogen and wheat. This strategy minimizes cell loss by preventing pathogen invasion without extensive mesophyll cell death, thereby conserving cellular resources.

The characteristics of Lr14a, such as its inducible expression following infection by an avirulent isolate, predicted transmembrane helices, and the ability to induce cell death, are reminiscent of transcription activator-like effector (TALE)-activated executor resistance genes. These genes confer resistance to the bacterial pathogens of the genus *Xanthomonas* ^13,24^. Unlike the typical interaction between an *R* protein and its corresponding effector protein, TALEs bind to the promoter of executor *R* genes, activating their expression and triggering cell death. While this mechanism was initially identified in bacteria-plant interactions, increasing evidence suggests that fungi may also secrete TALE-like effectors that trigger *R* gene expression. *R* gene expression patterns dependent on avirulent pathotypes have been observed in response to other pathogens, including the barley powdery mildew pathogen *Blumeria graminis* f. sp. *hordei*, the sunflower downy mildew pathogen *Plasmopara halstedii*, and the rice blast pathogen *Magnaporthe oryzae* ^29–31^. Recently, two barley leaf rust resistance genes, *Rph3* and *Rph7*, were cloned, both showing avirulent pathotype induction, suggesting that they could function as executor genes^32,33^.

Our results indicate that *Lr14a* may also function as an executor *R* gene. This suggests that *P. triticina* might encode a TALE-like protein that directly activates *Lr14a* expression or indirectly regulates its expression by targeting other transcription factors. TALEs typically contain nuclear localization signals (NLSs) for transfer into the host nucleus, a DNA-binding domain consisting of 15–20 almost identical modules that interact with the host promoter, a transcription factor binding motif domain for interaction with transcription factors, and an activation domain^24^. These features could serve as a guide for identifying AvrLr14a.

Our expression data revealed that *Lr14a* was specifically induced by the avirulent isolate in mesophyll cells, while it triggered cell death in guard cells. This suggests that Lr14a-mediated calcium influx activates cell-to-cell communication, transmitting defense signals from mesophyll cells to guard cells where hypersensitive response (HR) is initiated and pathogen invasion prevented. Several studies in C3 plant species have demonstrated that signals from mesophyll cells are necessary to drive stomatal responses^34–37^. Recent research has shown that various mobile molecules - such as small proteins, peptides, RNAs, metabolites, and second messengers are involved in transmitting extracellular stimuli from sensing cells to target cells (39). For instance, sucrose produced in mesophyll cells reaches the guard cell apoplast via the transpiration stream, where it is imported into guard cells by H^+^/sucrose symporters, regulating stomatal responses^38–41^. Light-regulated transcription factor HY5, located in mesophyll cells, induces the expression of mesophyll-derived peptides, which are secreted into the apoplast and recognized by receptors on stomatal precursors, ultimately controlling stomatal development^42,43^. Auxin similarly regulates stomatal development by repressing mesophyll-derived peptides^44^.

Ca²⁺, a key second messenger in plant immunity, plays a pivotal role in cell-to-cell signal transduction. The ROS/Ca²⁺ wave is considered a major mediator of signal transduction between cells and tissues^45^. Several Ca²⁺ channels, such as glutamate receptor-like channels (GLRs) and the tonoplast-localized TWO-PORE CHANNEL 1 (TPC1), have been reported to mediate long-distance signal transduction and initiate calcium-based plant defense signaling^46–48^. Furthermore, prolonged Ca²⁺ influx can activate metacaspases (MCs), which cleave PROPEPs to release Peps into the apoplast, where they are sensed by neighboring cells^49^. Based on these findings, we hypothesize that: Lr14a-mediated Ca²⁺ signaling may promote the generation of immune-related signaling molecules, reactive oxygen species (ROS), microRNAs (miRNAs), transcription factors and peptides, delivering them to guard cells to initiate HR. Further studies should focus on identifying the linker signaling molecules associated with Lr14a that may transmit defense signals from mesophyll cells to guard cells.

Distinct from the extensive host cell death typically triggered by other *R* genes, the guard cell-specific death induced by Lr14a-mediated Ca²⁺ influx presents a unique *R* gene function. Our findings provide the first insight into how a race-specific *R* gene modulates stomatal behavior to confer resistance against a fungal pathogen.

## Methods

### GCaMP6m-based calcium measurement in HEK293 cells

HEK293 cells were cultured in DMEM medium containing 10% fetal bovine serum and 1% penicillin/streptomycin at 37℃ and 5% CO_2_. Cells were seeded on 8-well chambered cover glasses (Thermo scientific) and were transfected using PEI-Max. The coding sequences of target genes were cloned into *pGGHEK* vector carrying 3xflag tag. Calcium imaging was performed after 24 hours post-transfection using confocal laser scanning microscope (CLSM-Leica Stellaris 5) equipped with a 40x oil objective (NA = 1.30). Cytosolic calcium levels were monitored using the Ca²⁺ indicator protein GCaMP6m, which was transiently co-transfected with the target genes. Protein accumulation was detected by western blot. The cells were stimulated with 5 mM Ca^2+^ in calcium imaging solution (1 mM CaCl_2_, 7.2 mM KCl, 107 mM NaCl, 1.2 mM MgCl_2_, 11.5 mM glucose, and 20 mM HEPES-NaOH, pH 7.2).

### Calcium influx measurement in *N. benthamiana*

The experiment was performed as described previously^9,19^. Briefly, *A. tumefaciens* strains carrying the target gene were infiltrated into transgenic GCaMP3 *N. benthamiana.* The coding sequences of N-terminal HA-tagged target genes were cloned into *pER8* vector^50^, which contains an estradiol-inducible promoter. Protein expression was induced by spraying 100 μM β-estradiol and 0.001% [v/v] Silwet L-77 at 24-hour post infiltration. Leaf discs were harvested 3 hours after estradiol treatment, placed in a 96-well plate, and incubated in the dark for 1 hour. GCaMP3 fluorescence was then measured using a Tecan SPARK microplate reader. Fluorescence values were normalized to the untreated control (F/F_UT_), where F represents the measured fluorescence at a given time point and F_UT_ is the average fluorescence for uninfiltrated samples at the corresponding time points. Protein accumulation was confirmed by western blot analysis.

### Calcium flux measurement in wheat leaf using NMT

Calcium flux measurement was performed using Non-invasive Micro-test Technology (NMT)^51^. 10-day old wheat seedlings (Arina*LrFor*) were infected by avirulent *P. triticina* isolate 96209 and virulent *P. triticina* isolate 95037. After 4dpi, the epidermis was removed to expose the mesophyll cells, which were then incubated in bathing solution (0.1 mM CaCl_2_, 0.1 mM KCl, 0.1 mM MES, pH = 6) at room temperature overnight before measurement. The unstable data in the first ten minutes were removed during data analysis.

### Injection of oocytes and electrophysiological measurements

Electrophysiological measurements and protein localization in oocytes were conducted as previously described^17^. Briefly, oocytes were injected with 50 nL of cRNA at a concentration of ≥300 ng/µL for wild-type or mutant Lr14a-GFP fusions. For electrophysiology oocytes were injected with 50 nL of cRNA at a concentration of 450 ng/µl. Non-injected oocytes and oocytes injected with 50 nL of nuclease free water were used as control. The oocytes were then incubated in ND96 medium at 18 °C for three days to allow translation of the proteins before performing confocal microscopy or electrophysiological measurements.

### Water-soaking-like phenotype on *Nicotiana benthamiana*

*Agrobacterium tumefaciens* carrying an overexpression construct for the wild-type (WT) or EMS-induced loss-of-function mutant of *Lr14a* was used to transiently transform *Nicotiana benthamiana* as previously described^13^. LaCl₃ and GdCl₃ was infiltrated 2 hour post Agrobacteria infiltration in a concentration of 2 mM, dissolved in water. Water was used for infiltration on control leaf spots. Leaves were scanned 3 days post inoculation using a Fusion FX Imaging System (Vilber Lourmat, Eberhardzell, Germany).

### Leaf rust infection

Leaf rust infection was conducted as previously described^13^. Briefly, leaf rust spores were stored at -80 °C. They were thawed at 42 °C for 1 min, mixed with FC-43 oil, and then sprayed with a high-pressure air sprayer on humid plants, 10 days after sowing at two-leaf stage. After the infection, plants were dried for 30 min and then covered with a plastic foil cover to maintain a high level of humidity and kept in the dark for 24 h.

### Determination of cell viability

Trypan blue staining was performed on *N. benthamiana* leaves to detect cell death, following a previously described protocol^52^. The staining solution (10 mL lactic acid, 10 mL phenol, 10 mL glycerol, 10 mL H₂O, and 40 mg trypan blue) was incubated in boiling water for 3 min. After this, the leaves and 40 mL of ethanol were simultaneously added to the staining solution and incubated for another 5 min until the green color completely disappeared. The leaves were transferred to the destaining solution (2.5 g/mL chloral hydrate) and incubated overnight with shaking.

3,3′-diaminobenzidine (DAB) was used for staining of reactive oxygen species to indicate a hypersensitive response^53^. Infected leaf segments were placed in the staining solution (10 mM ascorbic acid, 1 mg/mL DAB, pH = 3) under vacuum for 30 min, then incubated overnight in the dark. The leaf segments were subsequently transferred to a bleaching solution (ethanol: acetic acid: glycerol = 3:1:1) and incubated in boiling water for 15 min. Finally, the segments were stained in coomassie brilliant blue solution (0.2% coomassie brilliant blue R250 in 100% ethanol) for 1 min to visualize the uredia.

The epidermis and protoplast of *N. benthamiana* were stained by 40 μg/mL Fluorescein Diacetate for 5 min, while the intact leaves of *N. benthamiana* were stained for 30 min. 2 μg/mL PI were used to stain intact *N. benthamiana* leaves for 30 min. All the samples were kept in the dark during staining.

### qRT–PCR analysis

*Lr14a* expression in mesophyll cells and epidermal cells was detected by qRT–PCR. The peeled epidermis and isolated protoplasts were shock-frozen in liquid nitrogen and stored at −80 °C. RNA extraction, and first-strand cDNA synthesis were performed as described previously^13^. qRT–PCR was performed as described^54^. The thermocycling conditions were 95°C for 20 s, followed by 40 cycles of 95°C for 3 s and 60°C for 20 s for all targets. A subsequent melting curve assessment was performed to exclude potential primer dimers. Relative quantities were calculated and normalized to the reference gene *ADP* to obtain calibrated normalized relative expression values using CFX Maestro software (Bio-Rad).

### Stomatal aperture measurement

10-days old wheat seedlings were infected with *P. triticina* isolate 96209 and 95037, followed by an overnight incubation in the dark. The abaxial epidermis was peeled from a 2 cm leaf segment taken from the leaf tip at 4 dpi, and images were captured immediately after peeling. The stomatal apertures were measured using ImageJ.

### Particle bombardment

Particle bombardment of wheat epidermal cells was performed as described previously^55^. The coding sequence of *Lr14a* was cloned into the *pTA22* vector carrying maize ubiquitin 1 promotor. Leaf segments of primary leaves from 8-day-old plants of the wheat cultivar Bobwhite S26 were co-bombarded with 7 μg of *Lr14a* and 5 μg of *mRFP*. *mRFP* was cloned into the *pTA22* vector and used as reporter gene to indicate gene transformation. Leaf segments were incubated at 20°C with 80% relative humidity for 72 hours before taking image using confocal microscope CLSM-Leica Stellaris 5.

### Statistical analysis

Data for quantification analyses are presented as mean ± SEM. All analyses were performed using GraphPad Prism 9.0. The two-tailed Student’s *t*-test and one-way ANOVA followed by Tukey’s test was used for two-group comparisons and multiple comparisons.

## Acknowledgements

We thank Keiko Yoshioka and Thomas DeFalco for providing *N. benthamiana* GCaMP3 seeds; and Christoph Ringli for providing the vector *pER8*. This project was supported by the University of Zurich, and grants 310030_204165 and 310030_212382 from the Swiss National Science Foundation to B.K. and C.Z, respectively.

## Author Contributions

L.Y. and B.K. conceived the project. L.Y. and L.W designed and performed calcium flux measurements in wheat. B.N. and U.L. designed and performed electrophysiological experiments in oocytes. G.H. generated constructs for oocyte expression. L.Y. and S.Z. performed calcium imaging in HEK293 cells. L.Y. and E.J. performed leaf-rust infection-related experiments. M.H. provided help with bioinformatics analysis. C.Z. provided supervision, as well as experimental and scientific expertise. L.Y., C.Z. and B.K. analyzed the data. L.Y. and B.K. wrote the manuscript. All authors commented and approved the manuscript.

## Data availability

All data supporting the findings of this study are available within the manuscript.

## Supplemental information

**Extended Data Fig. 1.**
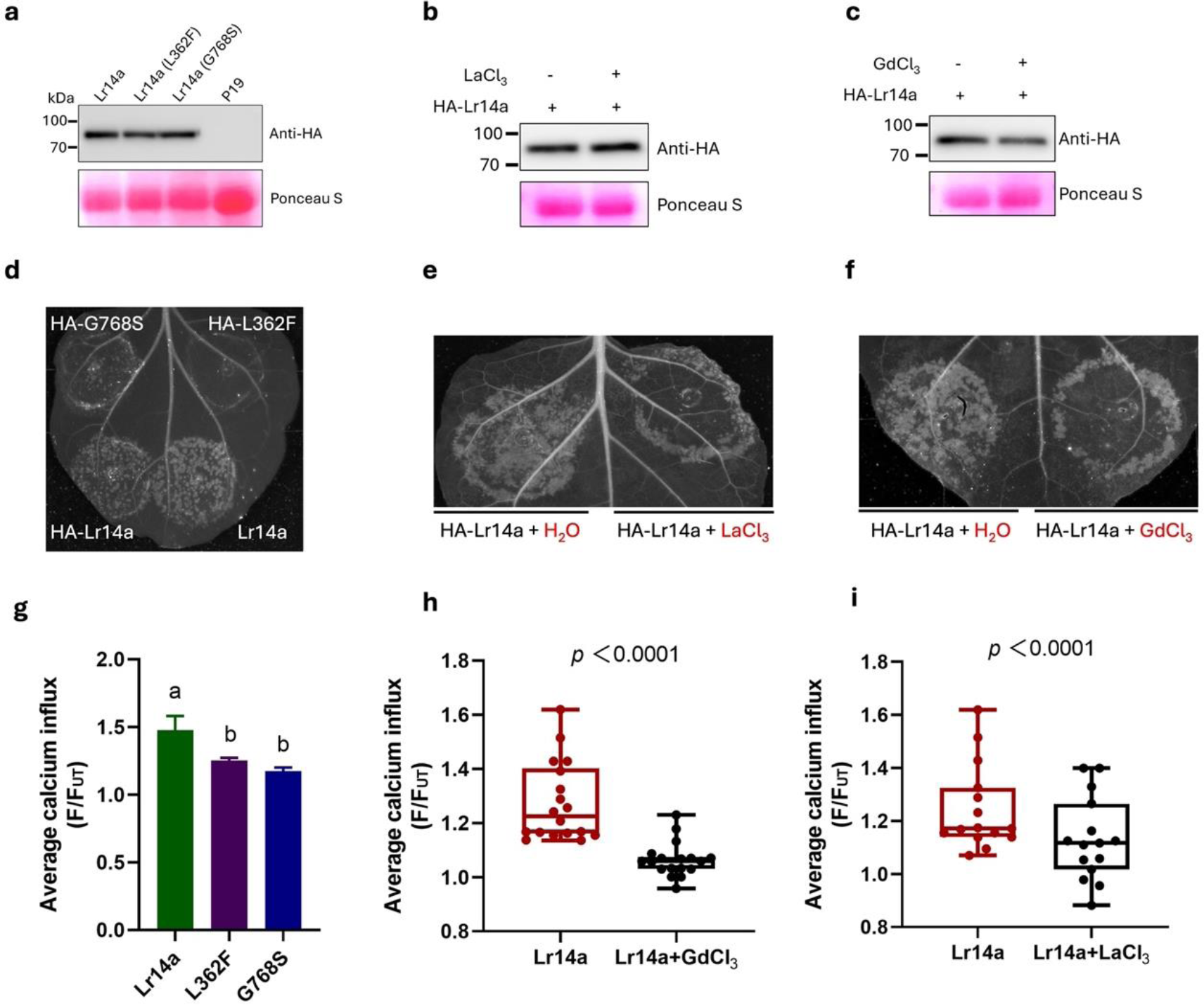
Protein accumulation and water-soaking-like phenotype caused by Lr14a and its mutants. **a-c**, HA-tagged *Lr14a* coding sequence was cloned into a *pER8* vector under the control of a β-estradiol inducible promotor. Protein accumulation was detected using western blot after 24-hour β-estradiol induction. *P19*-transformed leaves served as the negative control. Lr14a shows the same expression level with two mutants L362F and G768S (**a**). Protein accumulation was not influenced by the application of LaCl₃ (**b**) and GdCl₃ (**c**). **d-f**, water-soaking-like phenotype after expressing Lr14a or its mutants (**d**). The Lr14a-triggered water-soaking-like phenotype was abolished by the application of LaCl₃ (**e**) and GdCl₃ (**f**). **g,** Average calcium influx from all timepoint presented in Fig. 1a. Values are mean ± SE. n = 15-18 leaves, with different letters indicating statistical significance (*P* < 0.05). **h-i,** Average calcium influx from all timepoint presented in Fig. 1b (**h**) and Fig. 1c (**i**). Values are mean ± SE. Statistical significance was determined using an unpaired, two-tailed *t*-test.

**Extended Data Fig. 2.**
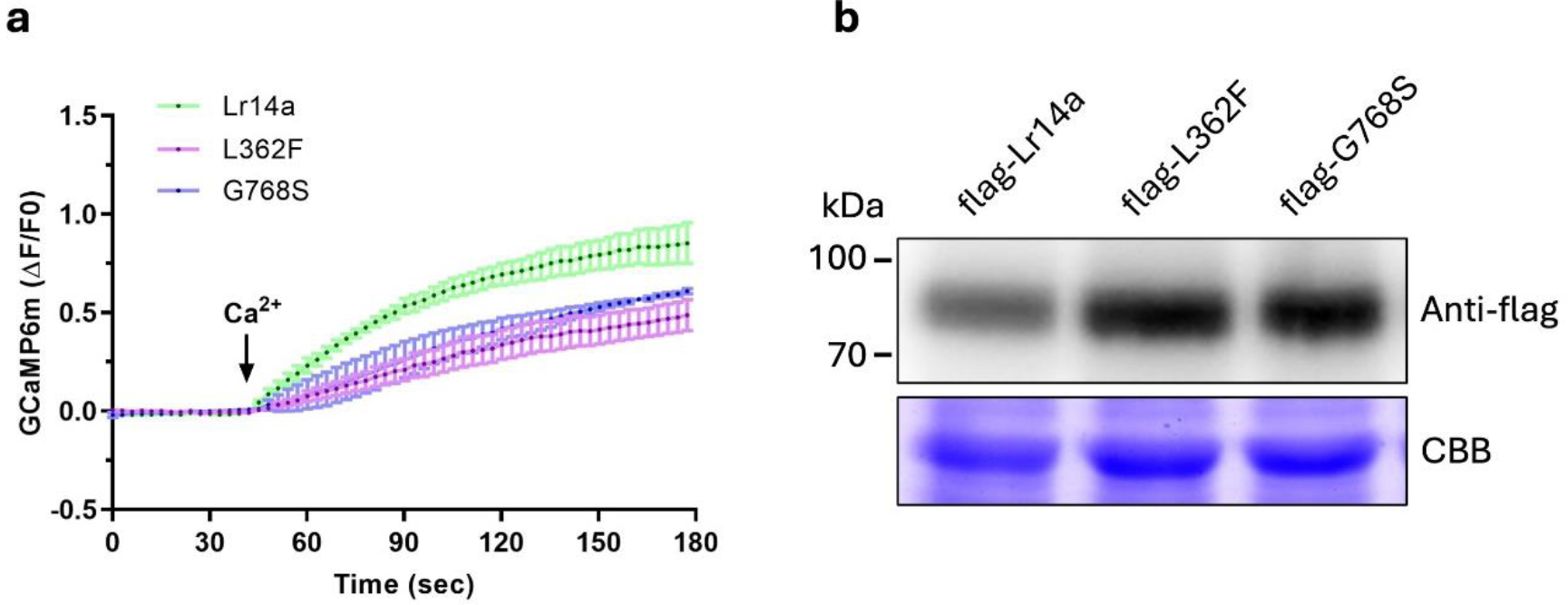
Lr14a-triggered [Ca^2+^]_cyt_ influx in HEK293 cell. **a**, Time-course recording of [Ca^2+^]_cyt_ influx in response to 5mM Ca^2+^ in HEK293 cells expressing flag-tagged Lr14a, or its mutants L362F and G768S. Changes in [Ca²⁺]_cyt_ were visualized using GCaMP6m. n = 3. **b**, Protein accumulation in HEK293 cells tested by western blot.

**Extended Data Fig. 3.**
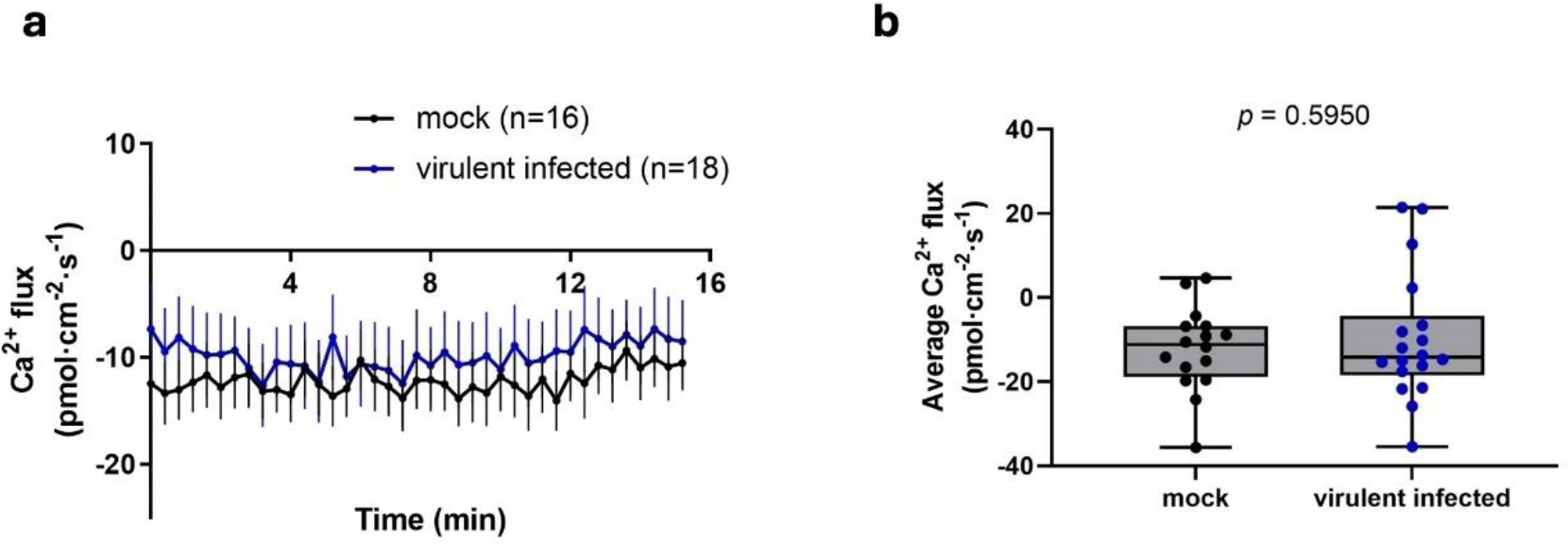
Ca^2+^ flux measurements in Arina*LrFor* after virulent *P.triticina* isolate 95037 infection at 4dpi. **a**, Time-course recordings of Ca^2+^ flux in mesophyll cells of Arina*LrFor* using the NMT technique. Negative values indicate Ca^2+^ influx, and positive values indicate Ca^2+^ efflux. **b**, Quantitative analysis of average Ca^2+^ fluxes from (**a**). The data are presented as mean ± SEM. Statistical significance was determined using an unpaired, two-tailed *t*-test.

**Extended Data Fig. 4.**
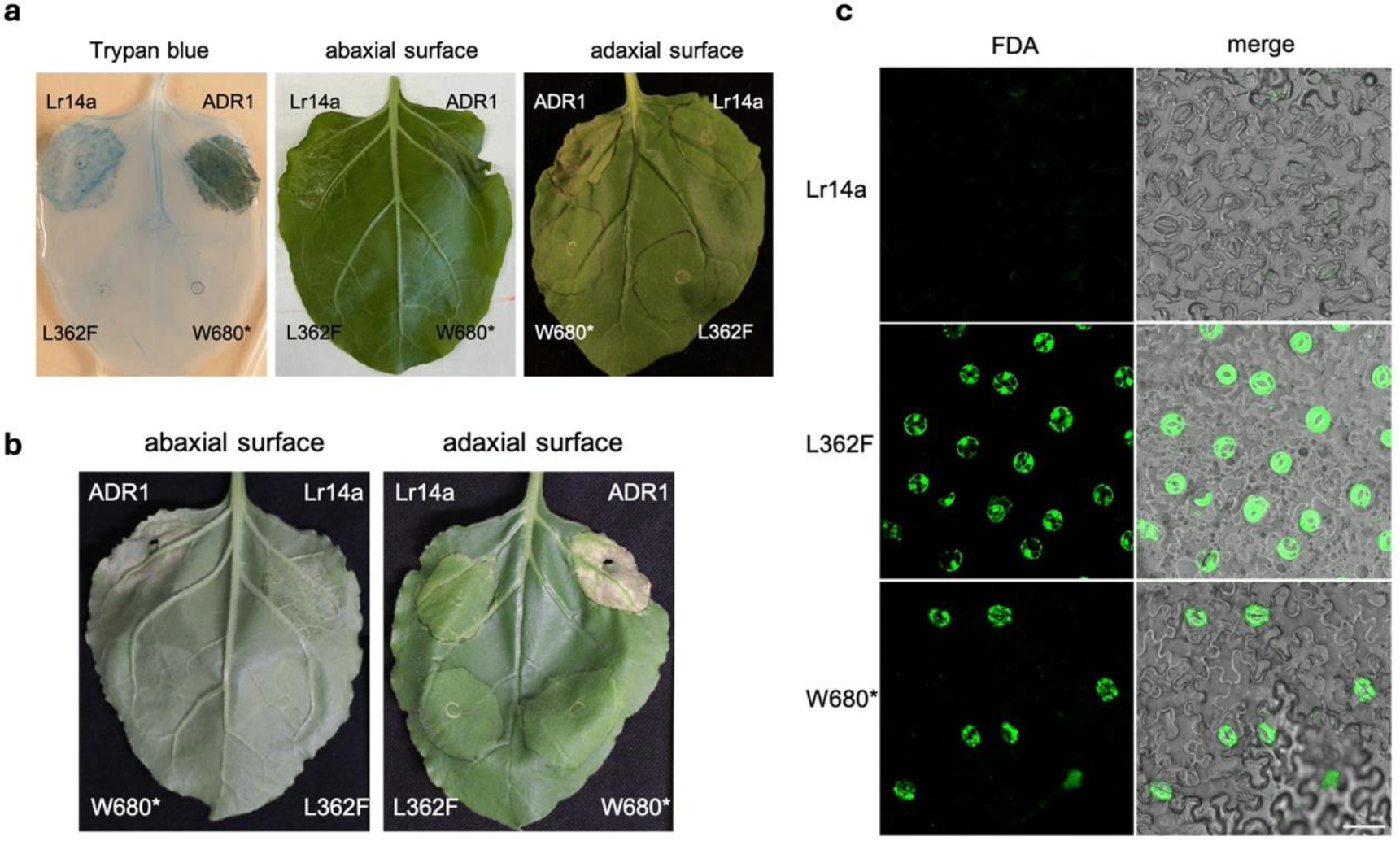
Lr14a induces guard cell death in *Nicotiana benthamiana*. **a**, Cell death and water-soaking-like phenotype in *N. benthamiana* leaves caused by the expression of HA-tagged wild-type (WT) Lr14a, autoactive ADR1, and two loss-of-function Lr14a mutants (L362F and W680*) at 3 days post-infiltration (dpi). Left: Representative image of *N. benthamiana* leaves stained with trypan blue. Middle: water-soaking-like phenotype on the abaxial leaf surface. Right: water-soaking-like phenotype on the adaxial leaf surface. **b**, Water-soaking-like phenotype. Abaxial (left) and adaxial (right) leaf surfaces of *N.benthamiana* after transiently expressing Lr14a and its mutants at 10 dpi. **c**, FDA staining of epidermal cells for 5min in *N. benthamiana* leaves at 3 dpi following expression of HA-tagged wild-type (WT) Lr14a and the two loss-of-function mutants Lr14a (L362F and W680*). Scale bar = 50 μm.

**Extended Data Fig. 5.**
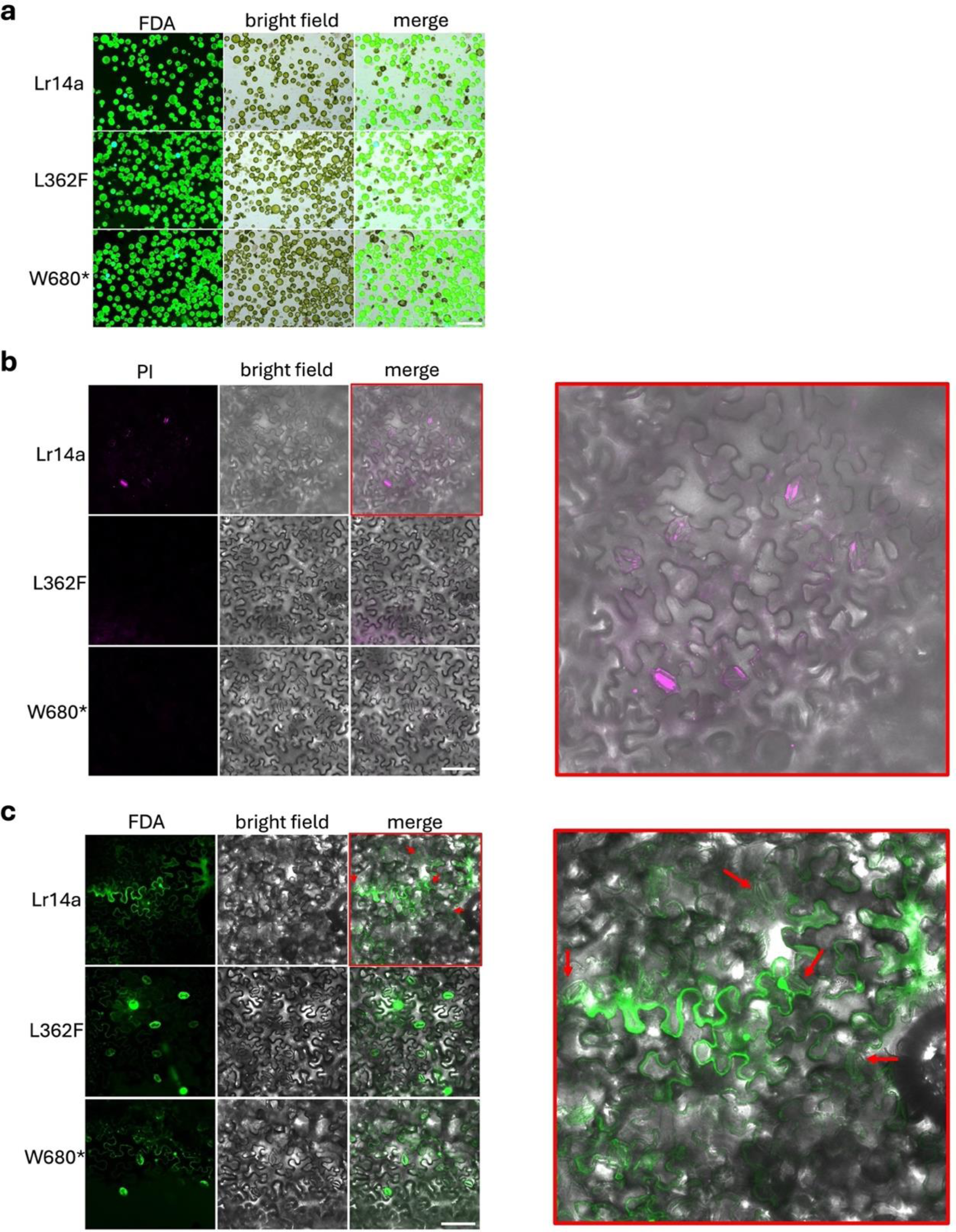
Lr14a triggers guard cell death in *N. benthamiana*. Cell viability was analyzed after transiently expressing *Lr14a* and its mutants at 3dpi using different dye. **a**, Protoplasts were isolated from leaves expressing Lr14a (top), L362F (middle), and W680* (bottom), and stained with FDA for 5 min at room temperature. Scale bar = 100 μm. **b**, Intact leaves expressing Lr14a (top), L362F (middle), and W680* (bottom) were stained with PI for 30 min at room temperature. The merged image of the Lr14a-expressing leaf was enlarged and marked with red boxes to show more detailed information. Scale bar = 100 μm. **c,** Intact leaves expressing Lr14a (top), L362F (middle), and W680* (bottom) were stained with FDA for 30 min at room temperature. The merged image of the Lr14a-expressing leaf was enlarged and marked with red boxes to show more detailed information. The wrinkled stomata were indicated with red arrows. Scale bar = 100 μm.

**Extended Data Fig. 6.**
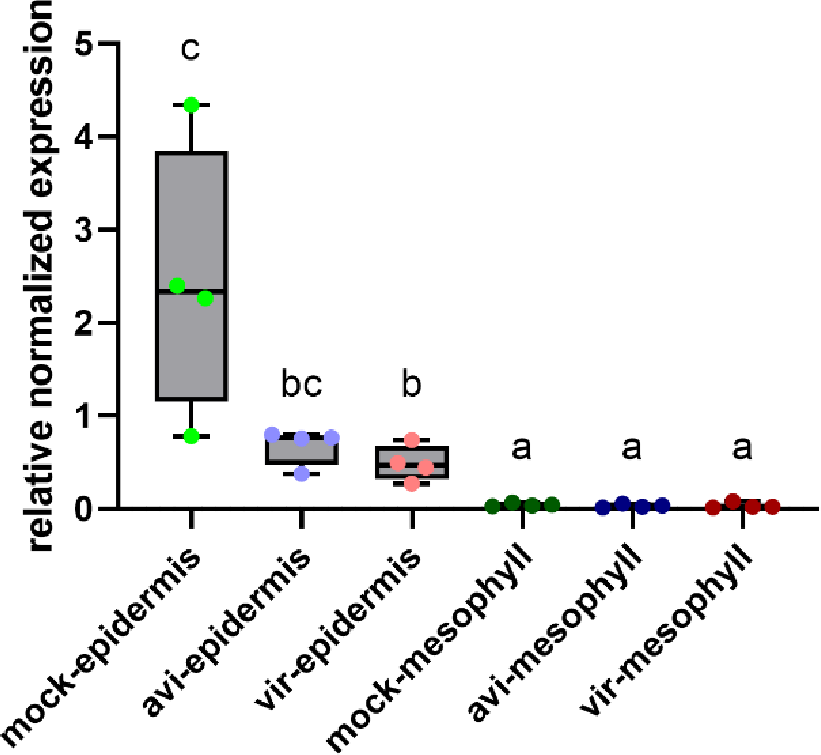
Expression of *TaALMT12* in mesophyll cells and epidermis. *TaALMT12* is a gene specifically expressed in guard cells. *TaALMT12* expression levels in peeled epidermis and isolated mesophyll cells were measured by qRT-PCR at 4 dpi after infection with avirulent isolate 96209 (avi) and virulent isolate 95037 (vir) isolates. n = 4, different letters indicate statistically significant differences (unpaired, two-tailed *t*-test).

